# Cellular Reprogramming in Bursts and Phases

**DOI:** 10.1101/2021.02.02.429448

**Authors:** Bradly Alicea

## Abstract

As a biochemical process, direct cellular reprogramming is slow and complex. The early stages of this process is the most critical determinant of successful phenotypic conversion. This study provides insight into the statistical signatures that describe temporal structure in the reprogramming process. We examine two sources of variation in reprogramming cells: clonal instances from various tissues of origin and rate of expansion between these lines. Our analytical strategy involved modeling the potential of populations to reprogram, and then applying statistical models to capture this potential in action. This two-fold approach utilizes both conventional and novel techniques that allow us to infer and confirm a host of properties that define the phenomenon. These results can be summarized in a number of ways, and essentially suggest that reprogramming is organized around changes in gene expression phenotype (phases) which happens sporadically across a cellular population (bursts).

## Introduction

The transition of cell phenotypes from pluripotent to differentiated states has traditionally been modeled as a series of state trajectories that undergo changes in phase. Two examples of this are the epigenetic landscape of Waddington (Ferrell, 2012) and the high-dimensional switch model of Cinquin and Demongeot (2005). These models have two assumptions in common. One assumption is that observed behavior can be characterized as a subset of all possible states (e.g. a phase space). The other assumption involves characterizing a transitory event as a bifurcation of a continuous function (e.g. the state trajectory). While these models are indeed suitable for describing differentiation as well as cellular reprogramming (the conversion from one cell phenotype to another), data from experiments on cell reprogramming suggest that a state space model does not fully describe variation in the process of phenotypic conversion. As a complement to the phase space approach, we propose a model of bursty dynamics that more fully characterize the variation observed both within and between instances of reprogramming populations (see Figure 1).

**Figure 1.**
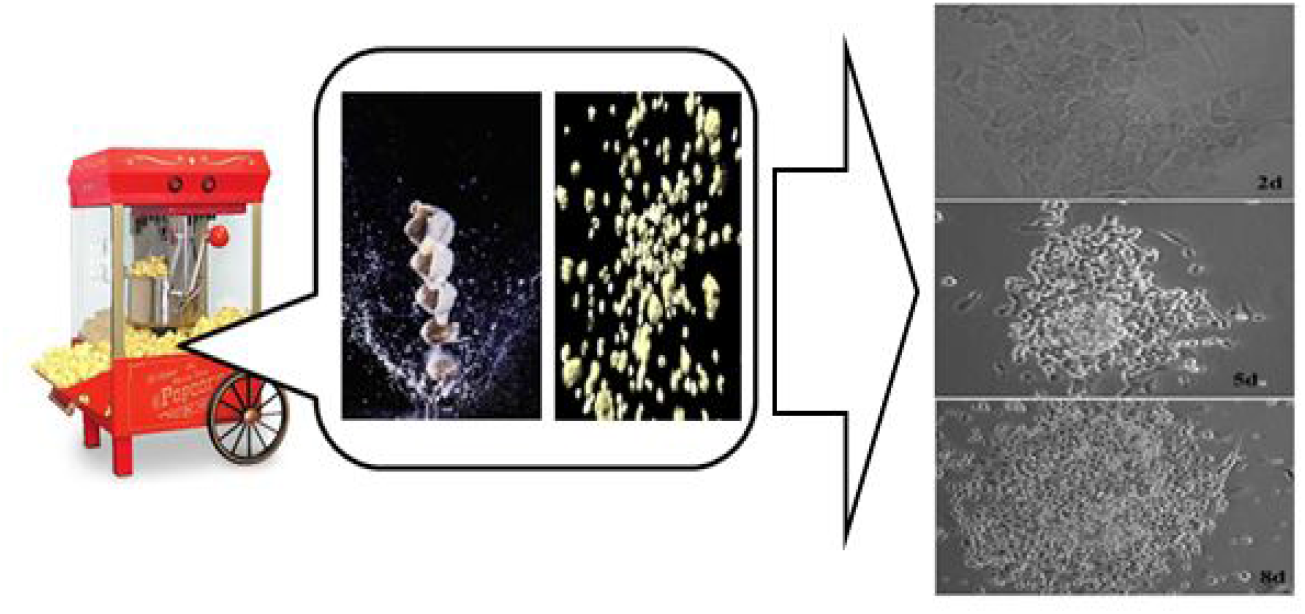
Cartoon depicting a physical analogue (popcorn) to approximate cellular reprogramming. Cells at right are pre-iPS cells at 2d (top), 5d (middle), and 8d (bottom) post-passage, and are used to illustrate colony formation.

From changes in phenotype (Tosh and Horb, 2004) to cascades related to gene expression and epigenetics (Richard et.al, 2011), cellular reprogramming can be defined as a series of large-scale effects arising from small-scale perturbations. One unresolved issue regarding the basic biology of direct reprogramming involves statistical signatures of variation that might help us better understand this process (Hanna et.al, 2010). The model is based on the idea that cellular reprogramming is a non-uniform process between and potentially within cell populations of different origins. To demonstrate this, it is shown that reprogramming events (defined using a time-aggregate measure of efficiency) can be characterized as a Poisson process. By using a rigorous measure of infectability, a rate of conversion can be established, which is shown to be variable among cell populations.

This state of affairs leads us to ask the following question: what sets up the reprogramming capacity of a given cell? To this end, we have modeled the growth and survival of multiple human and mouse cell lines using conditioned media and mathematical techniques to mimic the population-level phenomena that may affect cells during the earliest period of reprogramming (e.g. 2-8 days post-infection). Our primary goal is to employ a variety of analyses to establish that direct cellular reprogramming occurs in bursts and phases. It is these bursts and phases, defined in the next section, which places our unique datasets (see Methods) on infectability and reprogramming efficiency into context. To bridge the gap between bursts and phases, and to contextualize these bursts as characterizing changes in state, a phenomenological model called non-uniform gene regulatory networks will be introduced.

### Direct reprogramming as bursts and phases

In this paper, we will explore bursty and phasic behavior associated with direct cellular reprogramming. The bursty nature of cellular reprogramming can be understood by observing these events as variability in reprogramming efficiency. The proportion of cells in a population that convert to a new phenotype (e.g. reprogramming efficiency) varies from 0.001% to 29% for induced pluripotent stem cell (iPSC) reprogramming (Artyomov, Meissner, and Chakraborty, 2010). In the case of induced cardiomyocyte cells (iCMCs), the direct reprogramming approach of Ieda et.al (2010) yielded 15% of cells with cardiomyocyte-like expression patterns, but only 0.5% of cells became fully reprogrammed with the functional attributes of cardiomyocytes. A large range of efficiency values was also observed in the very first direct reprogramming experiments using MyoD cDNAs to induce cells to a skeletal muscle phenotype (Davis, Weintraub, and Lassar, 1987).

By contrast, bursty behavior of human (Barabasi, 2005) and physical (Karsai et.al, 2012) systems is generally understood as a series of short intervals of high activity interspersed among longer periods of low activity. In the context of direct reprogramming, bursty behavior can be defined as many-fold differences in reprogramming efficiency between cell lines. This can largely be inferred from counts over a finite time window within a single cell line, which allows for variation in reprogramming efficiency can be understood by using a Poisson (Barbour et.al, 1992) or some other exponential distribution.

Phasic behavior can be defined as a system that favors a particular physiological state in the face of environmental and other challenges to its identity (Izhikevich, 2007). These stable states have been proposed to exist during the process of cellular differentiation (Cinquin and Demongeot, 2005). Differences in these properties between cell lines are assumed to result from these lines undergoing conversion to a new phenotype at a variable rate (Hanna et.al, 2009; Yamanaka, 2009). Buganim et.al (2012) even proposes that cellular reprogramming occurs in two phases: an early stochastic phase in which transcriptional control is less hierarchical, and a later deterministic phase in which transcriptional control is more dependent on “master control” mechanisms. Yet phasic behavioral structure not only plays a role in the configuration of regulatory relationships, but behaviors of different cell populations in response to the reprogramming stimulus. As demonstrated in the case of phasic neurons, Gai, Doiron, and Rinzel (2010) observe a response invariant to small, low-frequency stimulation but an active response to large, high-frequency stimulation. In the context of direct reprogramming, phasic behavior can be defined as the proportion of cells that exhibit a signal for infection/transduction (e.g. YFP+) versus the proportion of cells that exhibit an additional signal for a cell type-specific marker (e.g. YFP^+^ and RFP^+^).

The discovery of signatures for bursty and phasic behaviors will be accomplished by using data from several independent experiments. Each experiment utilized a range of mouse fibroblast lines from various tissue types but all cloned from a single organism. By using count-based measurements of reprogramming efficiency and infectability, we are able to characterize variation across cell lines. This allowed us to directly count reprogramming events at two points in time and infer reprogramming events within a finite window of time. Comparisons could then be made within and between cell lines. Additionally, we were able to model population dynamics within a phase and between cell lines.

### Premises

To build an analytical strategy around the bursts and phases of direct cellular reprogramming, we will focus on three general premises: 1) cellular populations provide a “context” for reprogramming, 2) there are interesting relationships between population expansion and infectability, and 3) phenotypic conversions within an observable instance can be understood as a set of non-uniform events. Testing these premises will involve applying a Poisson model, curve-fitting to a power series, and a frequency analysis of rank-order behavior across replicates. To address the first premise, we utilize a strategy of growing each cell line in different types of conditioned media, and then using the resulting data to examine fluctuations in growth and survival between populations by introducing a technique called Population Fluctuation Analysis (PFA). The second premise can be addressed by looking at ways to predict infectability, and ultimately conversion. To accomplish this, we employ two types of analysis: a graphical representation of bivariate space (using bounding boxes) and variability of conversion rate. Investigating the third premise involves using statistical models based on an exponential distribution to understand the nature of direct reprogramming repeatability in terms of cell populations being exposed to a given concentration of virus.

### Methods

All molecular protocols and quantitative methods for cell segmentation and counting are fully described in Alicea et.al (2013). Quantitative data are freely available from Alicea et.al (2015), while all measures and datasets are described in the following subsections. All Appendices, Supplemental Figures, and Supplemental Tables can be found at https://doi.org/10.6084/m9.figshare.13644125.v1. Formal definitions of quantitative measures can be found in Appendix A, while more information about each dataset referenced in the paper can be found in Appendix B.

## Results

### Population Fluctuation Analysis (PFA)

A major barrier to our understanding of direct reprogramming efficiency involves fluctuation in cell number during exposure to a retrovirus and subsequent phenotypic remodeling. If a given cell population exhibits a high degree of cell death during this process, the overall efficiency is likely to be affected. One way to assess the effects of perturbation on different cell lines is to grow each cell lines under two sets of defined conditions (Dataset #1). The growth condition involves the use of growth-promoting media amenable to expansion of a population in the first days post-infection. The survival condition involves the use of serum-poor media, and is used to assess the contraction of a population in the first several days after initial exposure. Taken together, population expansion and contraction can be used to mimic selective forces imposed on a cell population during early direct reprogramming.

By taking the difference between per day growth rate and survival rates (parameter G_net_ shown in Equation 3), we were able to approximate potential fluctuations in population size during reprogramming. Simulating the dynamics of growth and survival over the 2d-8d time period provides a window into events that immediately precede and may directly influence reprogramming efficiency. It is also a more sophisticated modeling mechanism for understanding observed variation in infectability and reprogramming efficiency. Figure 3 shows the results of these simulations. In general, growth and survival of cells before conversion should act to constrain both infectability and reprogramming efficiency.

### PFA-based categories

There are four general trends across cell lines that provide a background for a typical population of a given cell type (Figure 2). The first category, represented by KI3, LI6, and KI2, is characterized by heavy death early (2d-4d) and slow growth later (5d-8d). The second category (KI5, LU3, TA6, TE4, LU6, and KI6) is characterized by near-baseline rates of population change early (2d-5d) and exponential growth later (6d-8d). The third category is represented by TE5 and SM1, and is represented by linear growth across the time course (2d-8d). The fourth category, represented by a single cell line (HE4), is characterized by linear growth early (2d-5d) with a saturation of growth late in the series (6d-8d).

**Figure 2.**
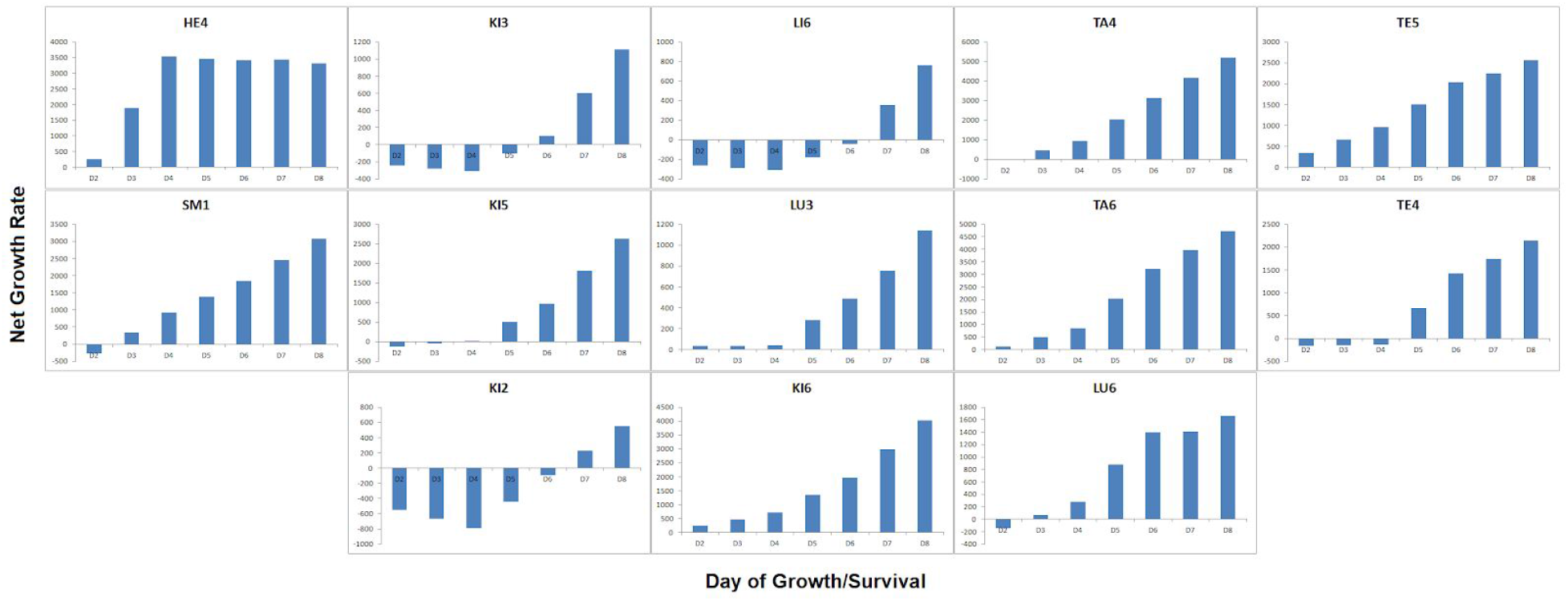
Simulations of growth and survival for all 13 mouse fibroblast cell lines from 2d to 8d. Net growth rate is measured as the number of cells on a particular day.

### Do these categories predict reprogramming?

Based on this model alone, we predict that the second group should exhibit the highest rates of reprogramming efficiency. Conversely, the first and fourth category should exhibit very low reprogramming efficiencies. When comparing this to reprogramming efficiency, the model is predictive of cell lines expected to exhibit low rates of reprogramming, but not those expected to exhibit high rates of reprogramming. This may be due to two factors: heavy death early reduces the population of cells that can be infected, while slow growth later restricts the number of cells that can result in full-blown converted phenotypes.

An alternate strategy is to look at those cell lines that exhibit high degrees of reprogramming efficiency and/or infectability. For reprogramming efficiency, TA6 and KI6 both exhibit exponential growth beginning at 2d. TA4 and KI5 also conform to a similar but less pronounced pattern. For infectability, TE4 and LU6 also exhibit exponential growth, but also exhibit the highest degrees of variation in the sample. LI6 is an exception to this, exhibiting heavy death throughout the time-course. However, there is a burst of growth late in this time course, which may lead to the proliferation of infected cells that will successfully convert to a new phenotype. On the other hand, KI3 exhibits a nearly identical pattern of population dynamics that do not reflect similar infectability or reprogramming efficiency statistics. This may point to an underlying mechanism present in LI6 cells but not in KI3 cells.

### Comparing PFA-based categorization with PCA

The question of how unique the population dynamics are for individual mouse fibroblast cell lines can also be addressed using a principle component analysis (PCA) of the growth and survival components. Figure 3 demonstrates that cell lines exhibiting variation along one the x- or y-axis (PC1 and PC2, respectively) should exhibit different behaviors in terms of either overall reprogramming efficiency or variation in reprogramming efficiency across type of viral infection or reprogramming variability within a cell line. An informal comparison of these analyses reveals that there may be some relationship between a population’s potential rate of population expansion, survival and reprogramming efficiency.

**Figure 3:**
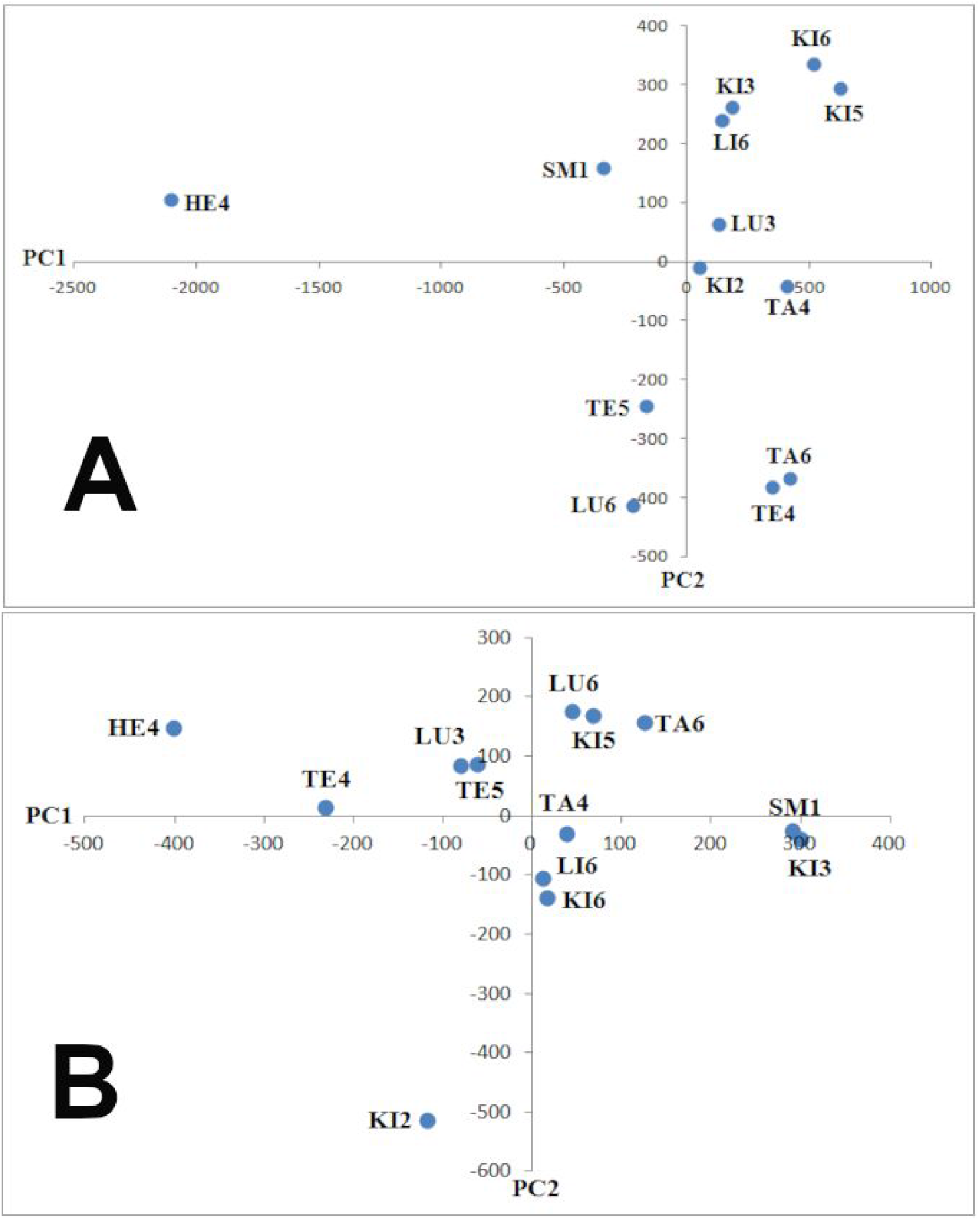
Principal component analysis (PCA) of cell counts for all cell lines under growth (A) and survival (B) conditions.

A PCA analysis on the growth component of our net growth rate measure revealed outliers along PC1 (HE4) and PC2 (TE5, LU6, TA6, TE4). While HE4 exhibited rapid expansion using the net growth rate measure, the result was split between linear expansion (TE5, TA6) and lagged expansion (LU6, TE4) for the outliers along PC2. The most extreme outliers along PC1 (HE4, TE4) and PC2 (KI2) for the survival component of our net growth rate measure exhibit very high and very low survival potential, respectively. When compared to a multidimensional scaling (MDS) analysis (Supplemental Figure 1) of the reprogramming efficiency for induced muscle (x-axis) and induced neuron (y-axis), we can see that the major outliers on the muscle dimension (KI6, TA4, KI5) show no relationship with the population fluctuation data. TA6, which exhibits linear expansion, is one of the more robust muscle converters. SM1 and KI3, which share variance along PC1 of the survival component, both share a lagged expression profile and slightly increased reprogramming efficiency for muscle.

### Do population dynamics predict infectability?

Aside from using an informal categorization technique, what else can population dynamics tell us about reprogramming? By examining differences between cells in terms of infectability, we can better understand how completely the viral vector penetrates the population prior to actual phenotypic conversion. Using an inducible YFP signal (Dataset #2), a rate of infectability can be assessed across cell lines. By assuming that the YFP and DAPI counts can yield independent rates, a derivative net growth rate can be established (shown in Equation 4). Figure 4 demonstrates these derivative net growth rates calculated across all cell lines for cells converting to neuron and muscle.

**Figure 4.**
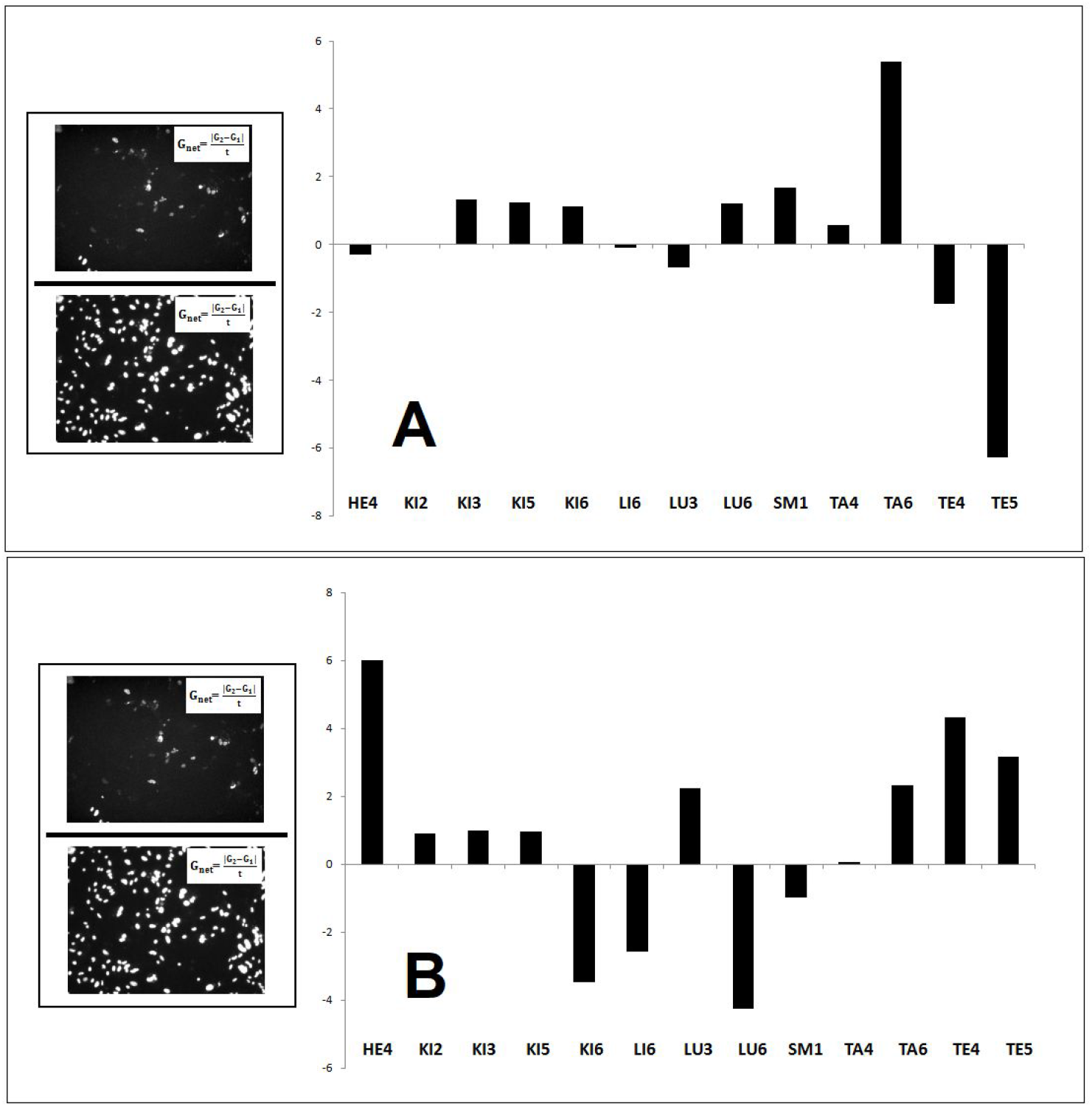
Derivatives of per day net growth rate (G_net_) for cells being actively converted to neurons (A) and muscle (B). Briefly, a derivative per day net growth rate is the G_net_ parameter calculated on YFP^+^ cell counts over the G_net_ parameter calculated on DAPI^+^ counts, where the YFP and DAPI counts are assumed independent. Images to the left of the bar graphs are from the same observation and are used to illustrate the derivative measurement of the YFP^+^ signal.

The purpose of examining the derivative per-day growth rate was to examine whether or not there were large changes in YFP^+^ cells independently of fluctuations in DAPI^+^ cells. In Figure 4, cell lines that exhibited large positive and large negative values were of particular interest. Large positive values represent a large-scale increase in the number of YFP^+^ cells relative to DAPI^+^ cells, while large negative values represent a large-scale decrease in the number of YFP^+^ cells relative to DAPI^+^ cells. For instances of cells actively converting to muscle, large increases are seen for HE4, TA6, TE4, and TE5. Large decreases are seen for lines KI6, LI6, and LU6. For instances of cells actively converting to muscle, large increases are seen for TA6. Large decreases are seen for both testes-derived lines (TE4 and TE5). Comparing these results to the PFA results (Figure 2), we can see that large increases in derivative G_net_ correspond with linear growth over time, while large decreases in derivative G_net_ often correspond to slight decreases in first-order G_net_ during the early phase of our time-series.

To assess the rate of conversion per hour, we can transform an aggregate count of YFP^+^ cells per observation into a per hour rate (see Figure 5). This rate is expected to be variable across observations for each cell line, and fall within a range of values that is variable across cell lines. A graph of conversion rate per hour using two types of measurements for cells actively converting to neurons and muscle is shown in Figure 6.

**Figure 5.**
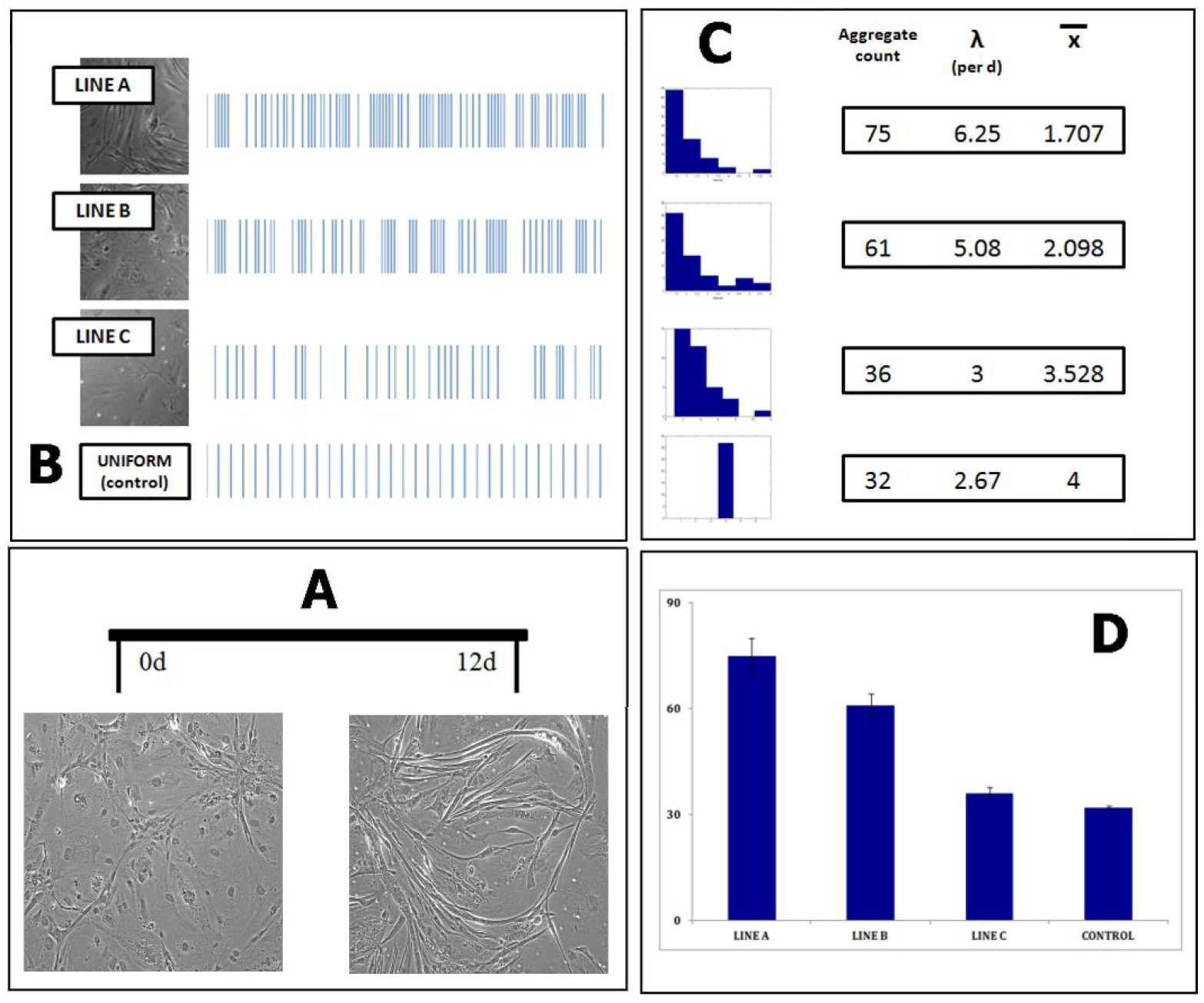
Diagrams showing the method by which counts for reprogramming over time are inferred. A: A picture of fibroblasts (0d, uninfected) converted to muscle fibers over a 12-day interval. B: Pseudo-data showing the expectation of “bursts” (e.g. non-uniformity over time) for different cell lines and comparison with a uniform control condition, C: Expected distribution of counts and statistics from pseudo-data in Frame B. D: Parametric summary of pseudo-data in Frames B and C, under the assumptions of a normal distribution.

**Figure 6.**
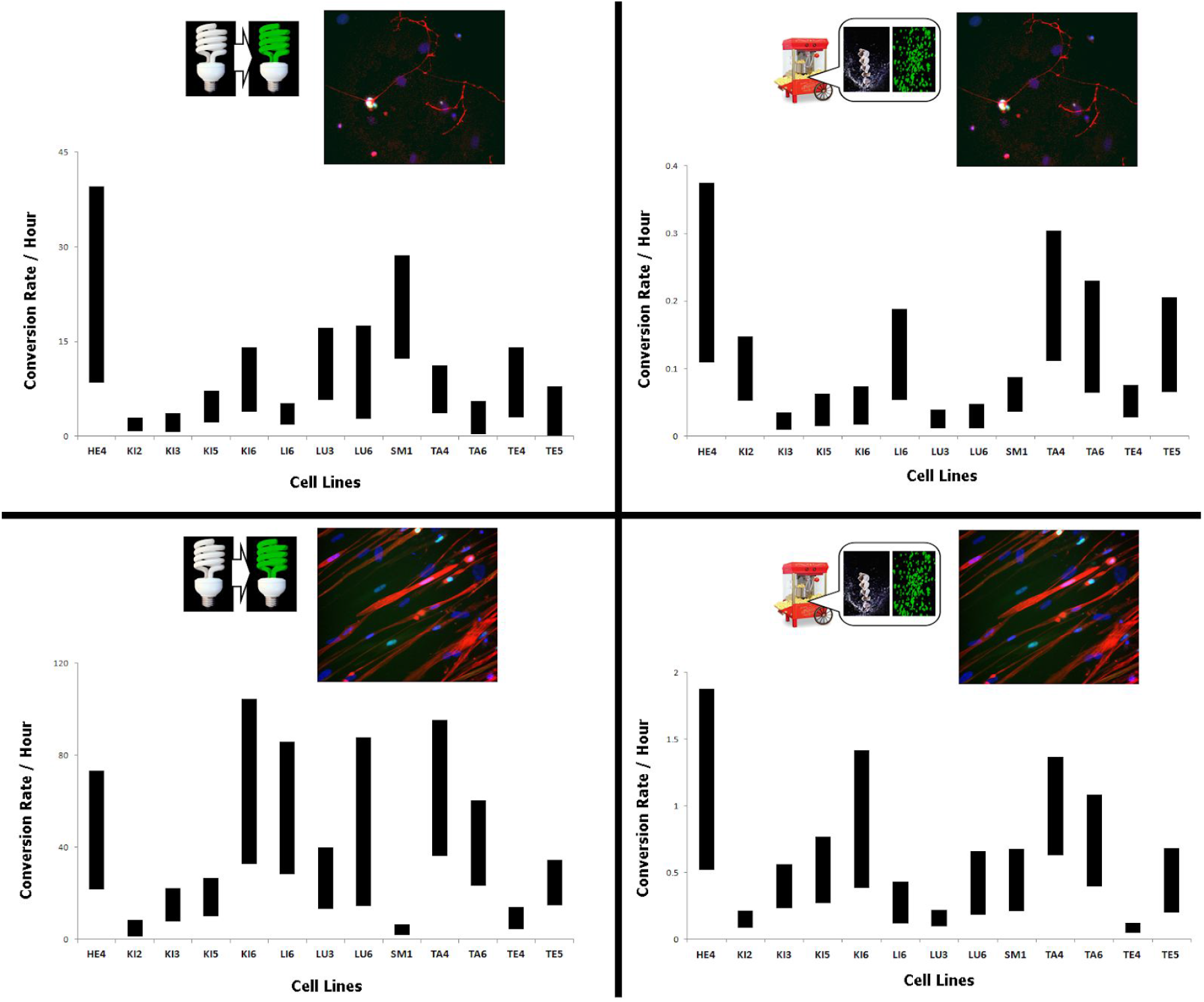
Variable rate analysis. Predicted rate of cell reprogramming, characterized as number of cells converted (conversion rate) per hour. **LEFT:** GFP-inducible model, **RIGHT:** actively converting cells. **TOP:** neuronal cells. **BOTTOM:** muscle cells.

Conversion rate per hour was assessed using both an YFP-inducible signal (Dataset #2) and YFP incorporated into a viral vector (Dataset #3). There are also two pieces of information that can be extracted from the plots in Figure 6: variability across instances of reprogramming and values for the rate itself. Taking both of these pieces of information into account, there are consistently high and low results. A high result characterized HE4 when converted to muscle, and HE4, KI6, and TA4 when converted to neuron. A low result characterizes KI3 and KI5 when converted to muscle, and TE4 and KI2 when converted to neuron. When comparing between the YFP-inducible and the YFP-viral conditions, a big difference is observed for lines TE5, TA6, LU3, and SM1 when converting to muscle, and for lines LI6, LU6, and SM1 when converting to neuron. The relationship between reprogramming efficiency (from Dataset #3) and mean net growth rate (per day, from Dataset #1) is restated by showing the range of variation in the form of bivariate bounding boxes (Figure 6).

Supplemental Figure 2 shows a comparison between infectability during a measurement of early infectability (4d) and later infectability (12d) using Dataset #4. These results demonstrate that the number of YFP^+^ cells are non-uniformly distributed within cell lines and exhibit many-fold differences between cell lines and over time. The count distributions (based on observed instances) are distinct when comparing early and later phases for the following cell lines: KI5, KI6, LU3, LU6, and TE4.

### Exponential models for within-cell lines diversity

The existence of a few cell lines with the ability to reprogram with much more facility than others is indicative of an underlying biological process that does not produce a normally-distributed outcome across biological diversity. This can be seen in Supplemental Table 1, where the fold-change compared to the population mean for each type of conversion has been calculated for each cell line.

When looking at population dynamics and calculating rates of infection and conversion, it becomes obvious that the response cell lines which would conventionally be referred to as outliers contain important information about the reprogramming process. In addition, the analysis of variability in the conversion rate (Figure 7) suggests that not all observed instances of reprogramming within a single cell line unfold in a predictable, deterministic manner. We argue that this type of diversity is to be expected, and should be characterized accordingly. To further explore this aspect of our data, we conducted three analyses more suited to understanding the relative frequency and magnitude of reprogramming-related events: an exact statistical test, calculation of rank order statistics, and curve-fitting to a power function.

**Figure 7.**
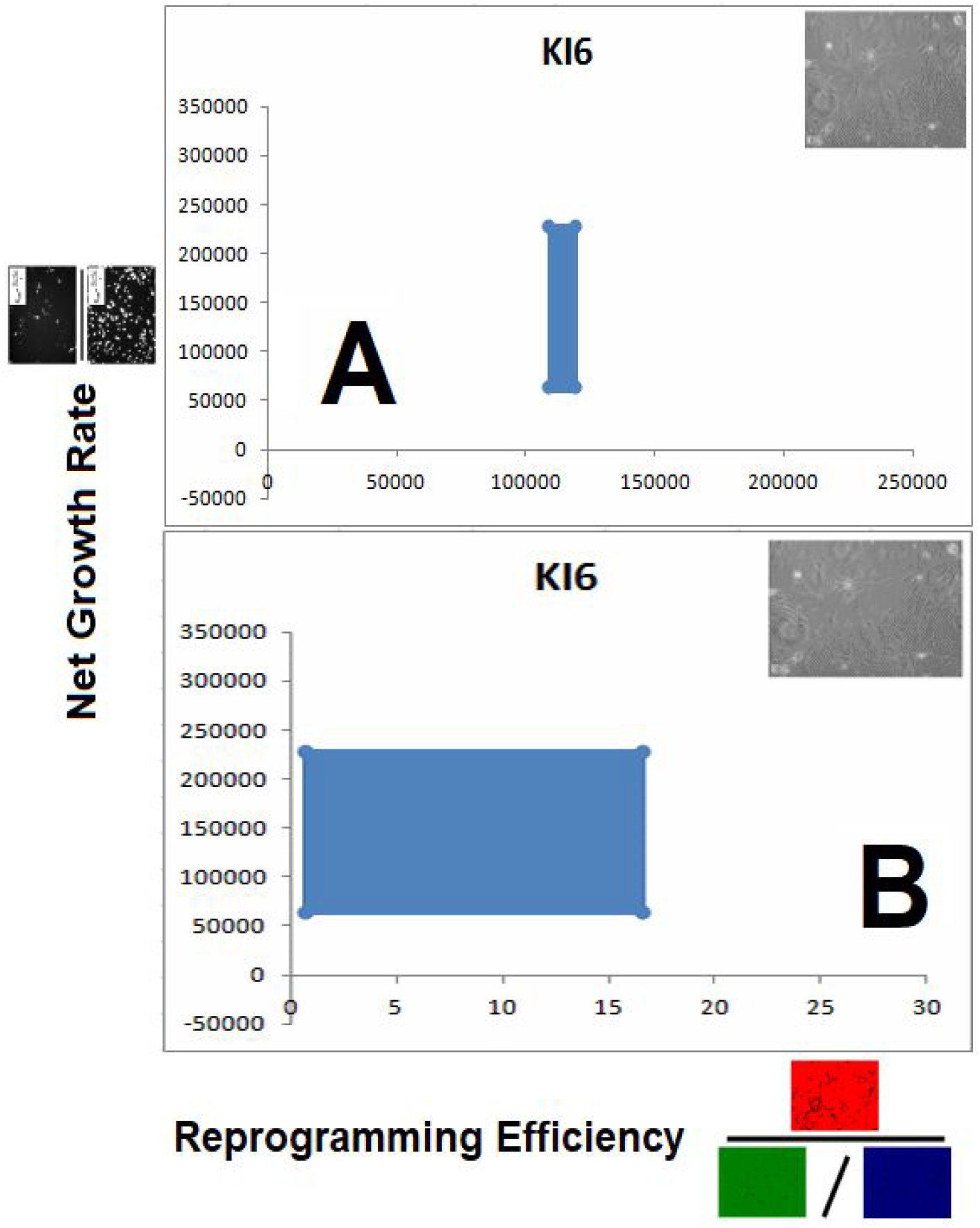
Example of the bounding box analysis for a selected cell line (KI6). The light blue box defines the range of observed reprogramming efficiency (RE) and per-day G_net_ values. Results are shown for muscle (left) and neuron (right).

Exact test. To assess the non-uniformity of reprogramming over time, a Poisson significance (exact) test was conducted on all cell lines for conversion to muscle (Figure 8) and neuron (Figure 9), the results of which are shown in Supplemental Table 2. Non-normality exists when a large proportion of samples fall outside a range of values determined uniformly about the mean. We can see that reprogramming efficiency exhibits more statistical significance for non-normality in neurons than in muscle. In neurons, 8 of the 13 lines sampled exhibited a non-normal distribution. For muscle, 5 cell lines (KI5, KI6, LI6, LU6, and TA4) exhibit non-normality. For infectability, the test for non-normality is largely not significant. Exceptions include TE4 and TE5 being converted to neuron and KI6 being converted to muscle.

**Figure 8.**
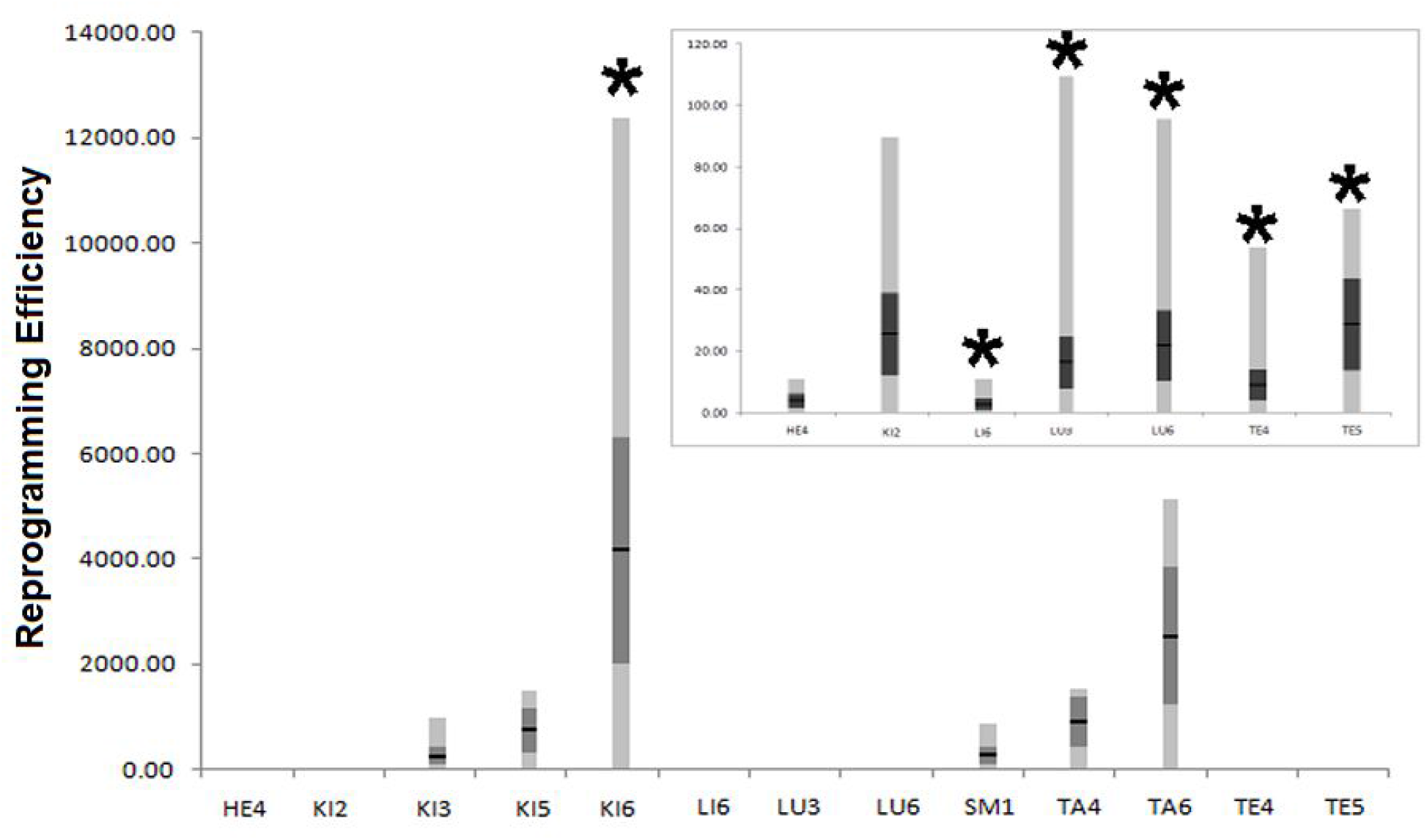
Possion (exact) test and distribution for all cell lines converted to muscle. Inset shows cell lines with small, non-zero values for reprogramming efficiency. Bars marked with a (*) are statistically significant at the .05 level.

**Figure 9.**
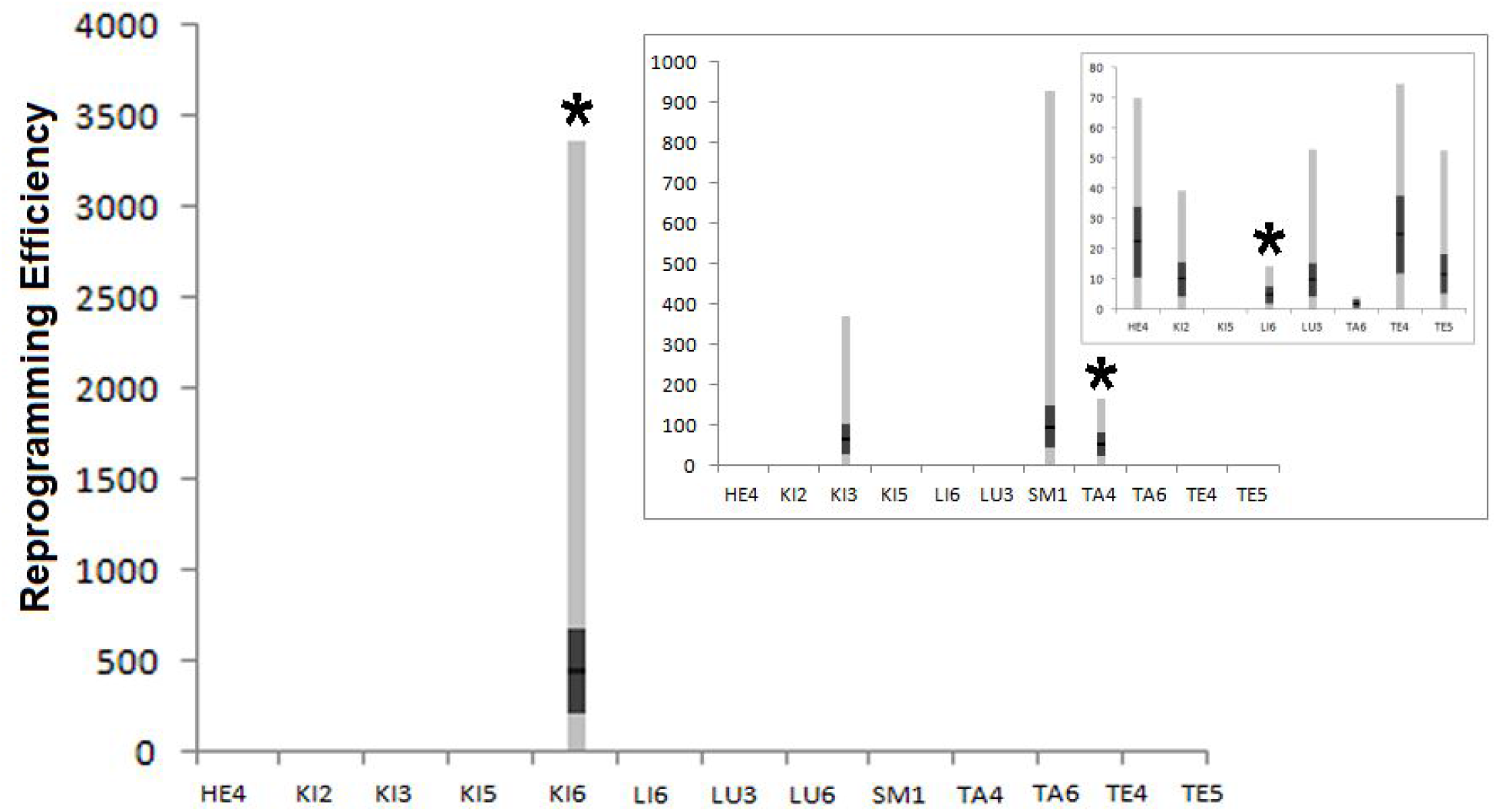
Possion (exact) test and distribution for all cell lines converted to neurons. Insets show cell lines with small, non-zero values for reprogramming efficiency at two different scales. Bars marked with a (*) are statistically significant at the .05 level.

Curve-fitting (power function). To further establish the statistical signature of large-scale differences in reprogramming over time, we used a power function to fit the data for each replicate involving conversion to muscle and neuron. Data for both reprogramming efficiency and infectability are shown in Supplemental Table 3. We calculated two parameters for these data: goodness-of-fit (R^2^) and power exponent (α). For the reprogramming efficiency measure, conversion to a neuronal phenotype fits the power function better as assessed by a higher overall R^2^ value. However, conversion to muscle results in a stronger power law. In terms of the 9 human cell lines (Dataset #5), conversion to neuron results in a result similar to mouse. Conversely, conversion to muscle results in an α-value much lower than for the mouse along with an overall lower R^2^ value.

### Rank-order frequency analysis

The second type of within-cell line analysis involves the calculation of rank order statistics and related parameters. Rank order frequency was calculated by sorting the reprogramming efficiency values for each replicate, and then calculating the frequency of a given rank position across all replicates for each cell line. Supplemental Tables 4 and 5 show the rank order frequency for muscle and neuron in the mouse, respectively. Supplemental Tables 6 and 7 show the rank order frequency for muscle and neuron in the mouse, respectively. Exemplar cell lines for the Dataset #3 (actively reprogramming mouse fibroblasts) are shown in Supplemental Figure 2.

We predict that if a given cell line is consistently ranked high (close to 1^rst^) or low (close to 13^th^) across a series of reprogramming instances, some innate mechanism is likely governing the outcome. Based on this criterion, all six mouse cell lines (KI6, TE5, TA4, HE4, LI6, and SM1) were selected as exemplars. Supplemental Figure 3 shows the histograms for both mouse (Dataset #3) and human (Dataset #5) cell lines after conversion to muscle and neuron. From this, we can see that mouse line KI6 is almost always 1^rst^ or 2^nd^, while mouse line HE4 is almost always 12^th^ or 13 ^th^ in muscle. In addition, different behaviors for the same cell line are seen when comparing conversion to muscle and neuron. The histogram for mouse line LI6 is bimodal in muscle, while for neurons the histogram is unimodal and spread across positions. The bimodal signature was ubiquitous for human cell lines converted to muscle. For human cell lines (Supplemental Figure 4 and Dataset 5) converted to neurons, a variety of results were seen. For example, human cell line 2 was mostly high ranking, while cell lines 7 and 9 were mostly low ranking. Cell line 4 exhibited a normal distribution around a median value of 5. These results may reflect the diverse genetic origins of the human cell lines.

## Discussion

To investigate our initial claims, we assess the stochastic fluctuations in the reprogramming process among different cell lines using PFA analysis. We then compare this method with a PCA analysis as a point of reference. In order to assess infectability, we then look at the population dynamics of the pre-programmed cells. This is done by comparing between muscle and neuron conversion in the same cell lines. For assessing non-uniform conversion, we assume that conversion conforms to a Poisson process, and deviates from a uniform process. Finally, a rank-order frequency analysis is used to assess consistent trends between cell lines. These provide additional clues to the normality, and perhaps more critically non-normality, of this process.

To better understand our results, we must make a distinction between individuality and diversity. In terms of building a phenotype, there are subtle differences between the two concepts. The concept of individuality among cells and organisms is based on the observation that cells within the larger can exhibit their own behaviors even when subject to a common developmental program (Wilson, 1999). In Buss (1987), individuality is treated as a common theme in development. One example of this individuality can be seen in the instability of the stem-like state in development. We would argue that individuality, which is driven by stochastic processes, may enable rare events such as many-fold increases in infectability and reprogramming efficiency. However, this individualistic response is not systematic across cell types, and can be buffered by normal developmental processes (Altshuler and Wu, 2010).

The study of bursts and phases can be applied to other biological processes, including cell division and differentiation in developmental systems such as Zebrafish and *C. elegans* (Alicea and Singh, 2021). In these systems, periodicity at multiple time scales have consequences on the spatial relationship between cells in a population. In the experiments shown here, cell populations are disembodied (conducted in cell culture). However, in an *in vitro context* (transplantation into an existing tissue), we might explore these bursty dynamics with respect to spatial organization.

## Supporting information

Supplemental Figure 1 (full resolution)

Supplemental Figure 2 (full resolution)

Supplemental Figure 3 (full resolution)

Supplemental Figure 4 (full resolution)

## Acknowledgements

I would like to thank Drs. Steven Suhr and Sarah Keaton for their technical expertise in data collection and cell culture. I would also like to thank the DevoWorm group for their feedback and discussions on how to measure burstiness in developmental systems.

