## Appendices and Supplemental Materials for "Cellular Reprogramming in Bursts and Phases"


| Section | Page |
|---|---|
| Appendix A: Quantitative Measures | 1 |
| Appendix B: Dataset Descriptions | 2 |
| Supplemental Figures | 3 |
| Supplemental Tables | 6 |


#### Appendix A: Quantitative Measures

Reprogramming efficiency (RE) was developed as a way to quantify all those cells or particles in an image that represent a successfully converted cell. Reprogramming efficiency can be defined by the following equation

$$RE = \frac{RED}{GFP:DAPI}^{-1} \qquad [1]$$

where RED represents the red channel, GFP represents the green channel and DAPI representes the blue channel in a *r,g,b* color scheme.

Infectability (I) was developed to quantify the proportion of cells that have successfully taken up the virus and expressed the transgenic elements related to cellular reprogramming. Infectability can be defined by the following equation

$$I = \frac{GFP}{DAPI} \qquad [2]$$

where GFP represents the green channel and DAPI represents the blue channel in a *r,g,b* color scheme.

Net Growth Rate. The population dynamics model consists of a series of net growth rate measurements, which consists of a comparison between the rates of growth and survival in idealized populations. This can be calculated using the following equation

$$G_{net} = \frac{|G_2 - G_1|}{t} - \frac{|S_2 - S_1|}{t} \qquad [3]$$

where $G_{net}$ is the net growth rate, $G_1$ and $G_2$ is the growth data sampled at two subsequent data points, $S_1$ and $S_2$ is the survival data sampled at two subsequent data points, and *t* is the time interval between the two points sampled.

Derivative Net Growth Rate. To understand the population fluctuation analysis in the context of cell infectability, the Gnet parameter can be calculated for both $YFP^+$ and $DAPI^+$ cell counts independently. This can be expressed as

$$G'_{net} = \frac{G_{net}(YFP)}{G_{net}(DAPI)} \qquad [4]$$

where $G_{net}(YFP)$ is the per-day net growth rate ($G_{net}$) for counts of $YFP^+$ cells, and $G_{net}(DAPI)$ is the per-day net growth rate ($G_{net}$) for counts of $DAPI^+$ cells

**Appendix B: Dataset Descriptions**

**Dataset #1.** Source fibroblasts are cultured under defined conditions, and then counted at four-day intervals. Defined conditions consisted of a growth media (serum-rich) and survival media (serum-deprived). Cells were cultured in each type of media for 12 days, which yielded a growth time-series and a survival time-series. Counts were interpolated to derive rates of growth and survival per day.

**Dataset #2.** YFP protein was localized to the nucleus of the source fibroblasts. These cells were then grown under the same defined conditions as described for Dataset #1. Cells are counted before and after plating, and this is done over three replicates. These counts were interpolated to derived rates of growth and survival.

**Dataset #3.** Source fibroblasts are actively reprogrammed using reprogramming factors delivered via retroviral vectors. Conversion to muscle fiber assessed at 13d, and conversion to neuron assessed at 30d. Immunostaining for cell type-specific markers (Tuj1 for neuron, alpha-actinin for muscle fiber) used to determine "red" channel signal. A flourescent (YFP) marker delivered to cells as part of the retroviral vector was used to determine infectability. These signals were normalized by a count of DAPI positive cells.

**Dataset #4.** Source fibroblasts are actively reprogrammed using reprogramming factors delivered via retroviral vectors. All viable and positive cells are counted at 4d and 12d using the immunostaining protocol described for Dataset #3.

**Dataset #5.** Human fibroblasts from the same tissue but a variety of genetic backgrounds were used to serve as a control dataset and place the source fibroblasts (single nude mouse) in context.

### Supplemental Figures

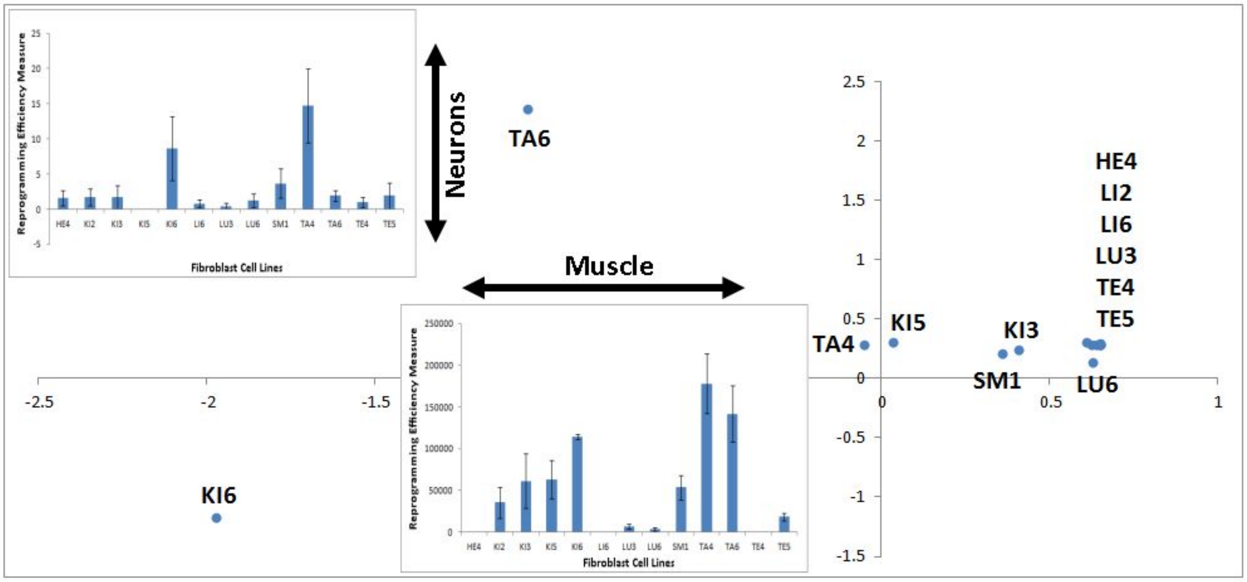


Supplemental Figure 1. MDS analysis for 13 mouse fibroblast cell lines reprogrammed to muscle (x-axis) and neuronal (y-axis) fates. Insets are a summary of descriptive results for neuronal reprogramming (upper left), and muscle reprogramming (lower center).

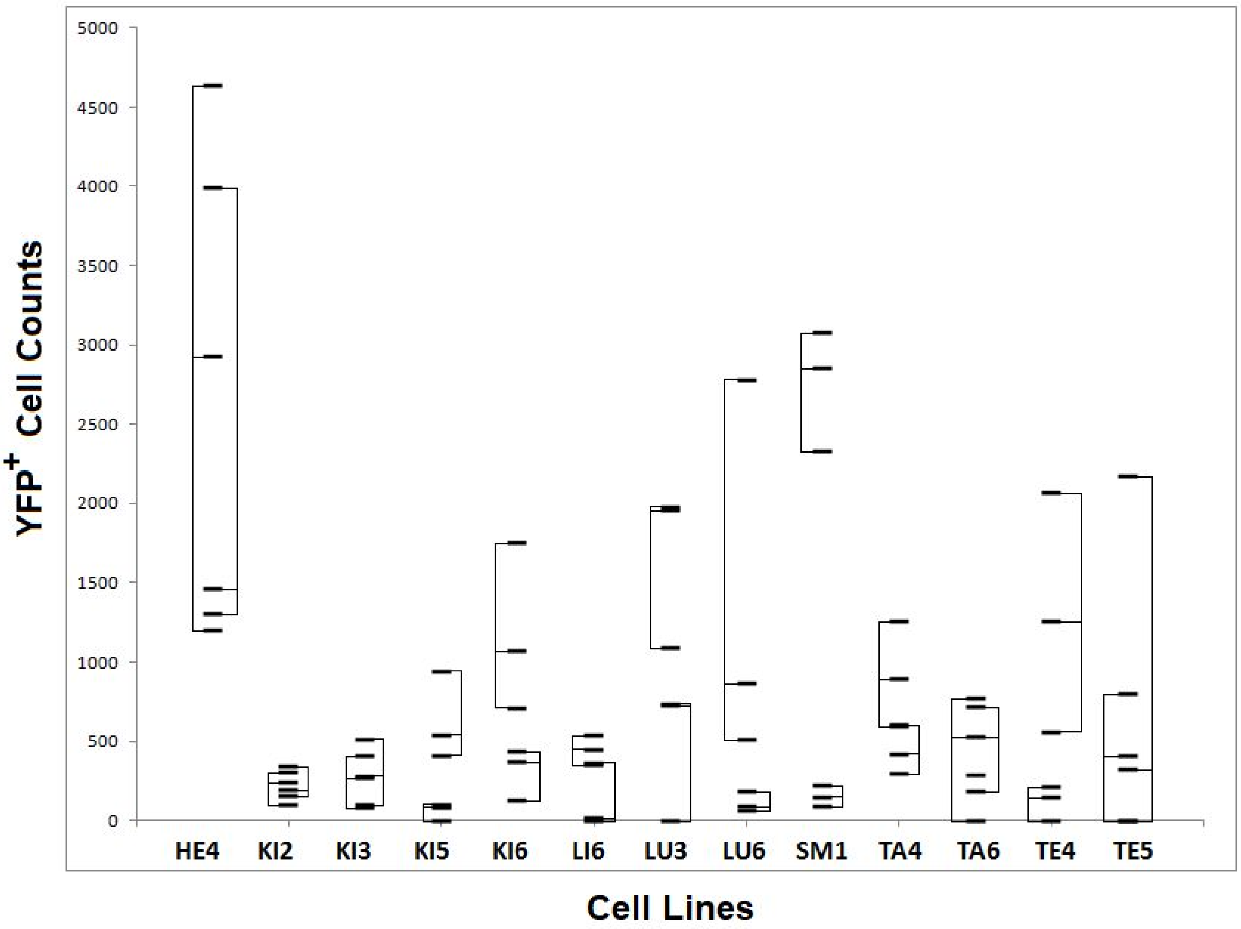


Supplemental Figure 2. Raw Counts of YFP$^{+}$ cells at two different phases of direct reprogramming (4d - early and 12d – later). Left-projecting bracket is the mean and range of the 4d values. Right-projecting bracket is the mean and range of the 12d values Cells being actively converted to muscle and neuron are treated as being part of the same condition.

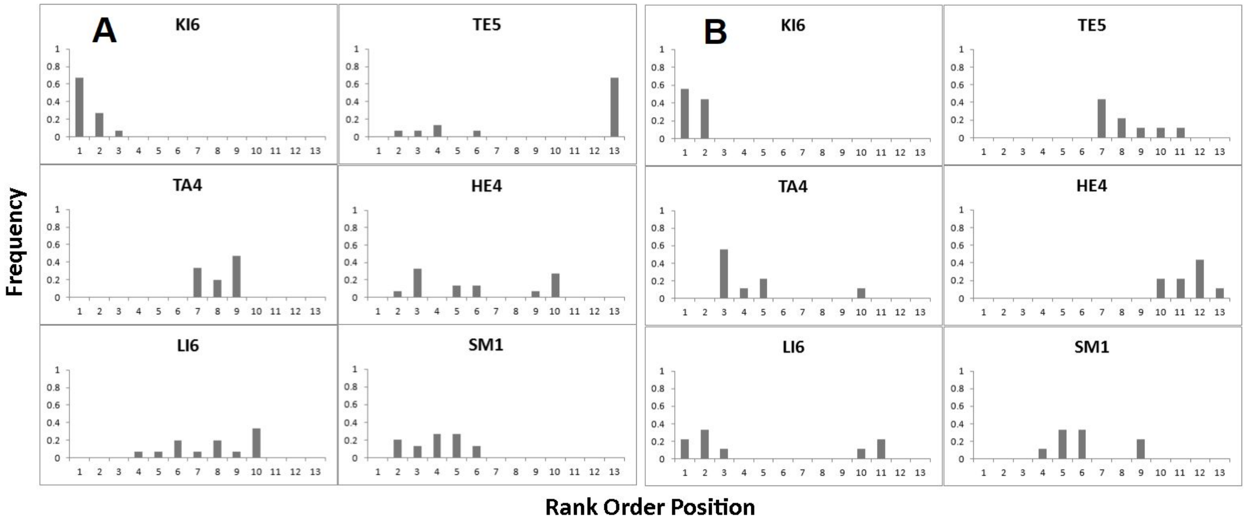


Supplemental Figure 3. Examples of the rank order histograms for six cell lines. LEFT: cell lines after conversion to neuron. RIGHT: cell lines after conversion to muscle.

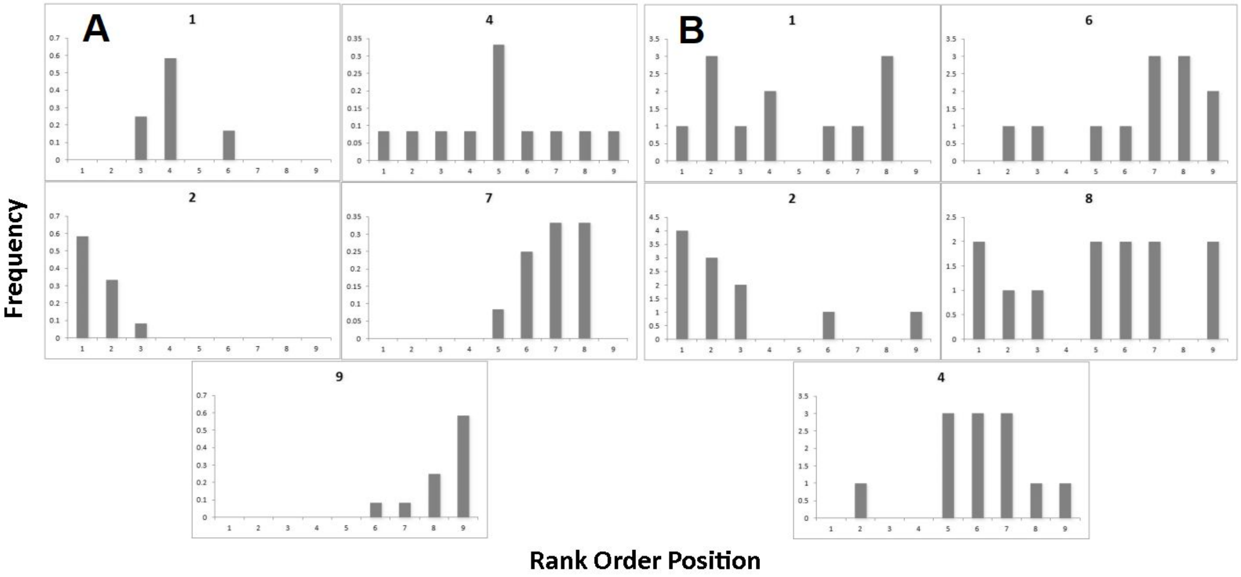


Supplemental Figure 4. Examples of the rank order histograms for six cell lines (from human dataset). LEFT: cell lines after conversion to neuron. RIGHT: cell lines after conversion to muscle.

**Supplemental Tables:**

Supplemental Table 1. Fold-change between global mean for all fibroblasts and specific lines.
* All instances were 0, comparison could not be made.

| | Reprogramming Efficiency (Muscle) | Reprogramming Efficiency (Neuron) | Infectability (Neuron) | Infectability (Muscle) |
|---|---|---|---|---|
| **HE4** | -179.34 | -2.57 | -1.92 | -1.60 |
| **KI2** | -26.94 | -5.74 | 1.31 | -2.86 |
| **KI3** | -2.70 | 1.10 | 1.13 | -1.83 |
| **KI5** | 1.07 | 0.00 * | 1.30 | -1.07 |
| **KI6** | 6.04 | 7.59 | -8.05 | -0.91 |
| **LI6** | -235.60 | -12.17 | 2.07 | 1.28 |
| **LU3** | -42.09 | -5.86 | -1.02 | -2.92 |
| **LU6** | -31.51 | 1.65 | -2.40 | 2.09 |
| **SM1** | -2.56 | -1.11 | -2.13 | -2.79 |
| **TA4** | 1.31 | -26.64 | 1.19 | -2.06 |
| **TA6** | 3.66 | -2.32 | 1.04 | -1.30 |
| **TE4** | -75.81 | -4.82 | 1.14 | 3.89 |
| **TE5** | -23.95 | -3.81 | 1.29 | -4.15 |

Supplemental Table 2. Poisson significance test for muscle and neuronal cell lines across mouse fibroblast cell lines. Each cell line and measure combination was resampled using 10,000 replicates (bootstrapping procedure). p-value < .05 is significantly different from normal distribution). Calculated using integers (rounded values). Each cell line taken independently.

| | **Muscle** | | **Neuron** | |
|---|---|---|---|---|
| **Cell line** | Reprogramming Efficiency | Infectability | Reprogramming Efficiency | Infectability |
| **HE4** | .48 | .63 | .00 | .83 |
| **KI2** | .08 | .21 | .00 | .24 |
| **KI3** | .20 | .15 | .67 | .54 |
| **KI5** | .00 | .14 | .53 | .79 |
| **KI6** | .02 | .00 | .05 | .69 |
| **LI6** | .00 | .09 | .00 | .33 |
| **LU3** | .31 | .28 | .00 | .43 |
| **LU6** | .08 | .58 | .00 | .10 |
| **SM1** | .45 | .70 | .62 | .23 |
| **TA4** | .00 | .28 | .53 | .55 |
| **TA6** | .53 | .32 | .11 | .93 |
| **TE4** | .32 | .23 | .00 | .04 |
| **TE5** | .51 | .21 | .00 | .00 |

Supplemental Table 3. Power law statistics for repetitions in cell lines converted to muscle and neuron for reprogramming efficiency (RE) and infectability (I). Parameters were estimated from linear-linear relationship.

| | Mouse Muscle (RE) | | Mouse Neuron (RE) | | Human Muscle (RE) | | Human Neuron (RE) | |
|---|---|---|---|---|---|---|---|---|
| Repetition number | $R^2$ | α | $R^2$ | α | $R^2$ | α | $R^2$ | α |
| 1 | .75 | 2.94 | .91 | 2.38 | .88 | 1.44 | 0.81 | 2.98 |
| 2 | .78 | 4.17 | .87 | 2.70 | .54 | 4.82 | 0.60 | 2.50 |
| 3 | .88 | 4.32 | .82 | 2.73 | .45 | 3.44 | 0.79 | 2.94 |
| 4 | .73 | 3.26 | .96 | 2.40 | .64 | 1.83 | 0.62 | 2.17 |
| 5 | .74 | 3.42 | .80 | 2.77 | .50 | 5.03 | 0.98 | 2.53 |
| 6 | .90 | 4.10 | .85 | 3.06 | .44 | 1.42 | 0.98 | 2.20 |
| 7 | .89 | 4.06 | .92 | 2.33 | .50 | 3.95 | 0.92 | 2.31 |
| 8 | .76 | 3.27 | .92 | 3.30 | .74 | 0.93 | 0.85 | 1.97 |
| 9 | .86 | 4.26 | .92 | 3.05 | .47 | 3.74 | 0.68 | 3.04 |
| 10 | - | - | .97 | 2.32 | .77 | 2.25 | 0.62 | 3.43 |
| 11 | - | - | .88 | 3.07 | .33 | 3.09 | 0.76 | 3.38 |
| 12 | - | - | .82 | 3.04 | .41 | 3.43 | 0.56 | 3.17 |
| 13 | - | - | .85 | 2.13 | - | - | - | - |
| 14 | - | - | .90 | 3.23 | - | - | - | - |
| 15 | - | - | .85 | 3.22 | - | - | - | - |

Supplemental Table 4. Consistency in rank order frequency domain across nine replicates for mouse cell lines after conversion to muscle. Each cell represents instances of position in rank order (high values equal consistency) across all replicates tested.

| | **Position in rank order (frequency of instances)** | | | | | | | | | | | | |
|---|---|---|---|---|---|---|---|---|---|---|---|---|---|
| | **1** | **2** | **3** | **4** | **5** | **6** | **7** | **8** | **9** | **10** | **11** | **12** | **13** |
| **HE4** | 0.00 | 0.00 | 0.00 | 0.00 | 0.00 | 0.00 | 0.00 | 0.00 | 0.00 | 0.22 | 0.22 | 0.44 | 0.11 |
| **KI2** | 0.00 | 0.00 | 0.00 | 0.00 | 0.00 | 0.11 | 0.22 | 0.33 | 0.00 | 0.33 | 0.00 | 0.00 | 0.00 |
| **KI3** | 0.00 | 0.00 | 0.00 | 0.22 | 0.33 | 0.44 | 0.00 | 0.00 | 0.00 | 0.00 | 0.00 | 0.00 | 0.00 |
| **KI5** | 0.00 | 0.11 | 0.33 | 0.44 | 0.11 | 0.00 | 0.00 | 0.00 | 0.00 | 0.00 | 0.00 | 0.00 | 0.00 |
| **KI6** | 0.56 | 0.44 | 0.00 | 0.00 | 0.00 | 0.00 | 0.00 | 0.00 | 0.00 | 0.00 | 0.00 | 0.00 | 0.00 |
| **LI6** | 0.22 | 0.33 | 0.11 | 0.00 | 0.00 | 0.00 | 0.00 | 0.00 | 0.00 | 0.11 | 0.22 | 0.00 | 0.00 |
| **LU3** | 0.00 | 0.00 | 0.00 | 0.00 | 0.00 | 0.11 | 0.22 | 0.00 | 0.00 | 0.11 | 0.11 | 0.11 | 0.33 |
| **LU6** | 0.00 | 0.00 | 0.00 | 0.00 | 0.00 | 0.00 | 0.11 | 0.33 | 0.44 | 0.00 | 0.00 | 0.11 | 0.00 |
| **SM1** | 0.00 | 0.00 | 0.00 | 0.11 | 0.33 | 0.33 | 0.00 | 0.00 | 0.22 | 0.00 | 0.00 | 0.00 | 0.00 |
| **TA4** | 0.00 | 0.00 | 0.56 | 0.11 | 0.22 | 0.00 | 0.00 | 0.00 | 0.00 | 0.11 | 0.00 | 0.00 | 0.00 |
| **TA6** | 0.44 | 0.44 | 0.11 | 0.00 | 0.00 | 0.00 | 0.00 | 0.00 | 0.00 | 0.00 | 0.00 | 0.00 | 0.00 |
| **TE4** | 0.00 | 0.00 | 0.00 | 0.00 | 0.00 | 0.00 | 0.11 | 0.11 | 0.11 | 0.00 | 0.11 | 0.11 | 0.44 |
| **TE5** | 0.00 | 0.00 | 0.00 | 0.00 | 0.00 | 0.00 | 0.44 | 0.22 | 0.11 | 0.11 | 0.11 | 0.00 | 0.00 |

Supplemental Table 5. Consistency in rank order frequency domain across nine replicates for mouse cell lines after conversion to neuron. Each cell represents instances of position in rank order (high values equal consistency) across all replicates tested.

| | **Position in rank order (frequency of instances)** | | | | | | | | | | | | |
|---|---|---|---|---|---|---|---|---|---|---|---|---|---|
| | **1** | **2** | **3** | **4** | **5** | **6** | **7** | **8** | **9** | **10** | **11** | **12** | **13** |
| **HE4** | 0.00 | 0.07 | 0.33 | 0.00 | 0.13 | 0.13 | 0.00 | 0.00 | 0.07 | 0.27 | 0.00 | 0.00 | 0.00 |
| **KI2** | 0.00 | 0.00 | 0.00 | 0.07 | 0.07 | 0.20 | 0.20 | 0.47 | 0.00 | 0.00 | 0.00 | 0.00 | 0.00 |
| **KI3** | 0.20 | 0.07 | 0.07 | 0.00 | 0.00 | 0.00 | 0.00 | 0.07 | 0.00 | 0.40 | 0.20 | 0.00 | 0.00 |
| **KI5** | 0.00 | 0.00 | 0.00 | 0.00 | 0.00 | 0.00 | 0.00 | 0.00 | 0.33 | 0.00 | 0.40 | 0.27 | 0.00 |
| **KI6** | 0.67 | 0.27 | 0.07 | 0.00 | 0.00 | 0.00 | 0.00 | 0.00 | 0.00 | 0.00 | 0.00 | 0.00 | 0.00 |
| **LI6** | 0.00 | 0.00 | 0.00 | 0.07 | 0.07 | 0.20 | 0.07 | 0.20 | 0.07 | 0.33 | 0.00 | 0.00 | 0.00 |
| **LU3** | 0.00 | 0.00 | 0.00 | 0.00 | 0.20 | 0.07 | 0.13 | 0.00 | 0.00 | 0.00 | 0.27 | 0.40 | 0.00 |
| **LU6** | 0.13 | 0.33 | 0.13 | 0.07 | 0.00 | 0.00 | 0.00 | 0.00 | 0.00 | 0.00 | 0.00 | 0.33 | 0.00 |
| **SM1** | 0.00 | 0.20 | 0.13 | 0.27 | 0.27 | 0.13 | 0.00 | 0.00 | 0.00 | 0.00 | 0.00 | 0.00 | 0.00 |
| **TA4** | 0.00 | 0.00 | 0.00 | 0.00 | 0.00 | 0.00 | 0.33 | 0.20 | 0.47 | 0.00 | 0.00 | 0.00 | 0.00 |
| **TA6** | 0.00 | 0.00 | 0.13 | 0.07 | 0.27 | 0.20 | 0.27 | 0.07 | 0.00 | 0.00 | 0.00 | 0.00 | 0.00 |
| **TE4** | 0.00 | 0.00 | 0.07 | 0.27 | 0.07 | 0.07 | 0.20 | 0.00 | 0.00 | 0.00 | 0.00 | 0.00 | 0.33 |
| **TE5** | 0.00 | 0.07 | 0.07 | 0.13 | 0.00 | 0.07 | 0.00 | 0.00 | 0.00 | 0.00 | 0.00 | 0.00 | 0.67 |

Supplemental Table 6. Consistency in rank order frequency domain across nine replicates for human cell lines after conversion to muscle. Each cell represents instances of position in rank order (high values equal consistency) across all replicates tested.

| | **Position in rank order (frequency of instances)** | | | | | | | | |
|---|---|---|---|---|---|---|---|---|---|
| | **1** | **2** | **3** | **4** | **5** | **6** | **7** | **8** | **9** |
| **1** | 0.17 | 0.33 | 0.17 | 0.08 | 0.08 | 0.08 | 0.08 | 0.00 | 0.00 |
| **2** | 0.42 | 0.25 | 0.08 | 0.17 | 0.08 | 0.00 | 0.00 | 0.00 | 0.00 |
| **3** | 0.17 | 0.17 | 0.42 | 0.08 | 0.17 | 0.00 | 0.00 | 0.00 | 0.00 |
| **4** | 0.00 | 0.00 | 0.08 | 0.17 | 0.17 | 0.25 | 0.08 | 0.17 | 0.00 |
| **5** | 0.00 | 0.00 | 0.00 | 0.08 | 0.08 | 0.17 | 0.33 | 0.17 | 0.08 |
| **6** | 0.00 | 0.00 | 0.00 | 0.00 | 0.17 | 0.17 | 0.00 | 0.25 | 0.42 |
| **7** | 0.00 | 0.00 | 0.00 | 0.08 | 0.08 | 0.00 | 0.33 | 0.33 | 0.17 |
| **8** | 0.17 | 0.08 | 0.00 | 0.17 | 0.08 | 0.17 | 0.08 | 0.00 | 0.25 |
| **9** | 0.08 | 0.08 | 0.08 | 0.17 | 0.17 | 0.08 | 0.08 | 0.08 | 0.08 |

Supplemental Table 7. Consistency in rank order frequency domain across nine replicates for human cell lines after conversion to neuron. Each cell represents instances of position in rank order (high values equal consistency) across all replicates tested.

| | **Position in rank order (frequency of instances)** | | | | | | | | |
|---|---|---|---|---|---|---|---|---|---|
| | **1** | **2** | **3** | **4** | **5** | **6** | **7** | **8** | **9** |
| **1** | 0.08 | 0.25 | 0.25 | 0.17 | 0.08 | 0.17 | 0.00 | 0.00 | 0.00 |
| **2** | 0.08 | 0.08 | 0.08 | 0.17 | 0.17 | 0.08 | 0.25 | 0.08 | 0.00 |
| **3** | 0.08 | 0.17 | 0.17 | 0.00 | 0.17 | 0.17 | 0.08 | 0.00 | 0.17 |
| **4** | 0.17 | 0.08 | 0.08 | 0.25 | 0.00 | 0.08 | 0.25 | 0.00 | 0.08 |
| **5** | 0.00 | 0.17 | 0.25 | 0.00 | 0.17 | 0.08 | 0.08 | 0.08 | 0.17 |
| **6** | 0.42 | 0.00 | 0.17 | 0.08 | 0.08 | 0.08 | 0.08 | 0.08 | 0.00 |
| **7** | 0.00 | 0.00 | 0.00 | 0.00 | 0.25 | 0.00 | 0.17 | 0.33 | 0.25 |
| **8** | 0.17 | 0.17 | 0.00 | 0.33 | 0.00 | 0.25 | 0.08 | 0.00 | 0.00 |
| **9** | 0.00 | 0.08 | 0.00 | 0.00 | 0.00 | 0.08 | 0.00 | 0.42 | 0.33 |
