## Supplementary figures and images for "Cellular Reprogramming in Bursts and Phases"

### Supplemental Figure 1 (full resolution)

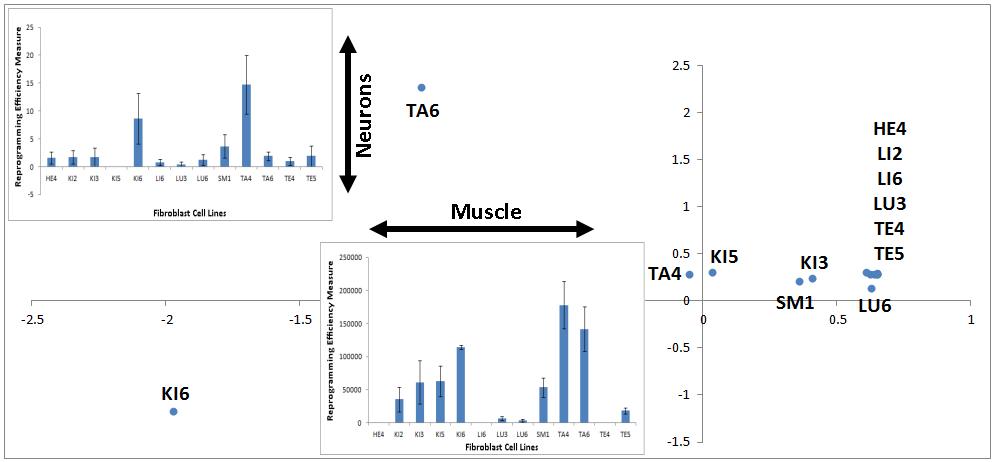

### Supplemental Figure 2 (full resolution)

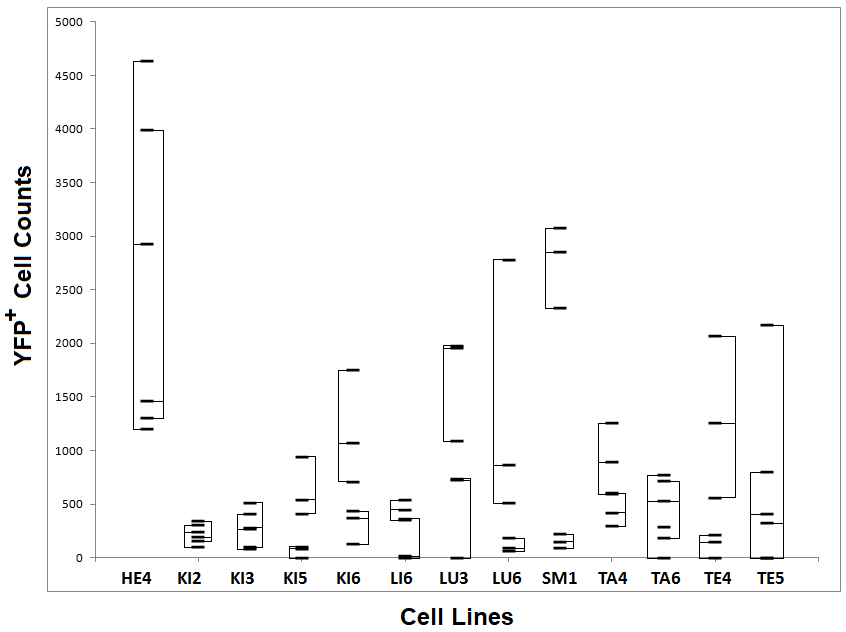

### Supplemental Figure 3 (full resolution)

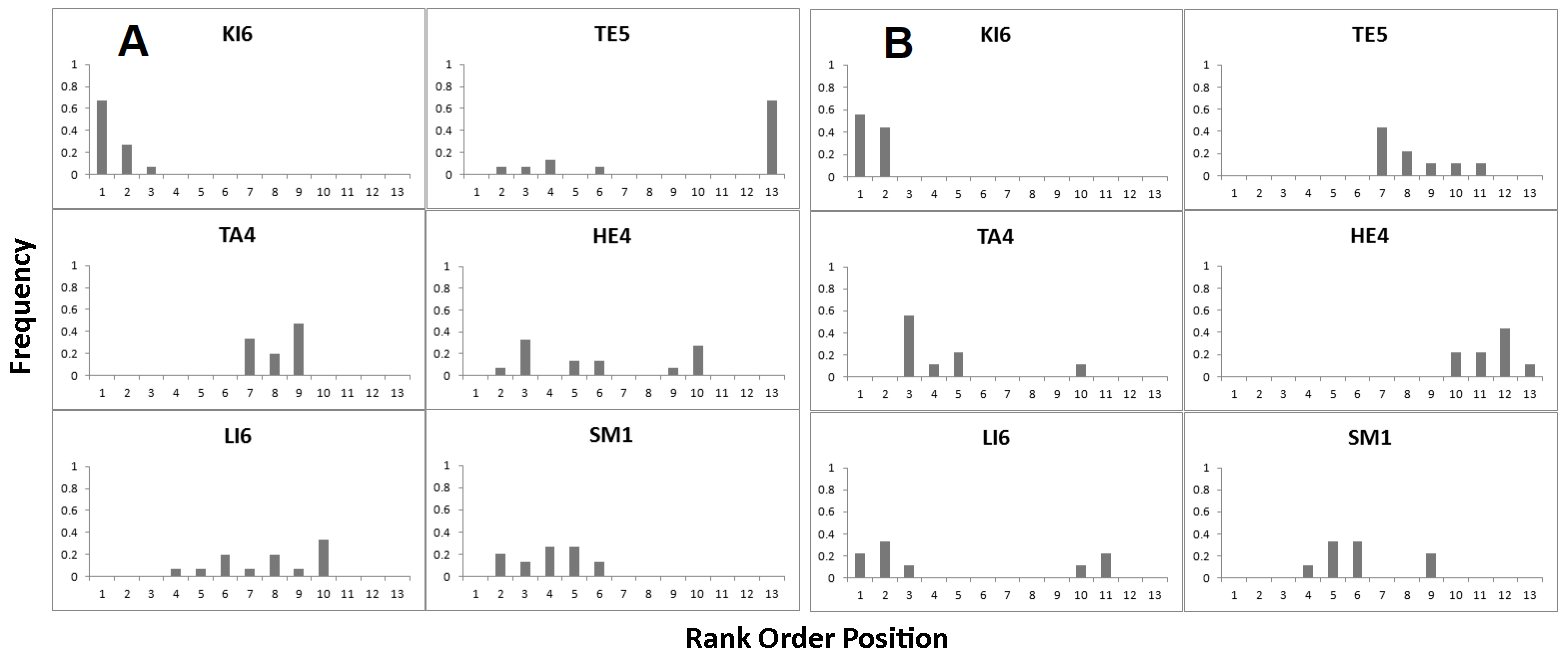

### Supplemental Figure 4 (full resolution)

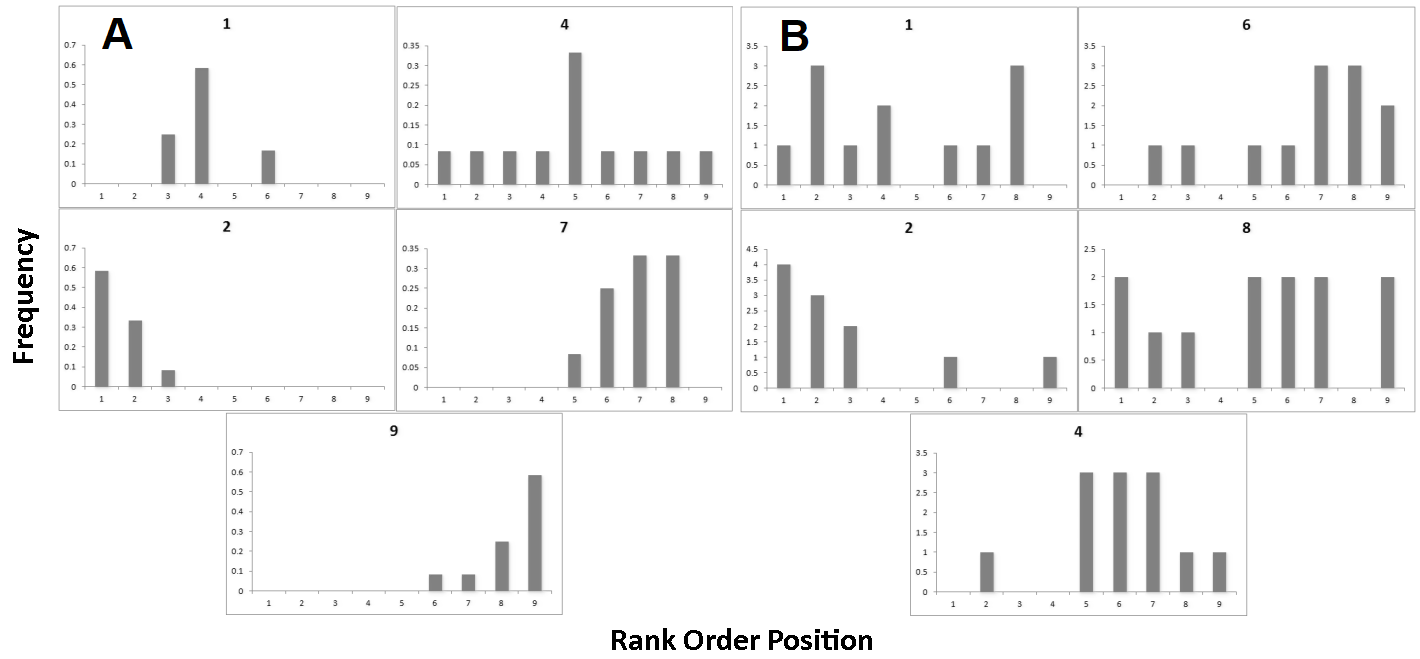
